# Late Integration of Prior Expectations During Precision-Weighted Perceptual Decisions

**DOI:** 10.64898/2026.04.13.718329

**Authors:** Timothy Gastrell, Dragan Rangelov, Jason B. Mattingley

**Affiliations:** Queensland Brain Institute The University of Queensland; Department of Psychological Sciences Swinburne University of Technology; School of Psychology The University of Queensland

## Abstract

Adaptive perceptual decision-making relies on the ability to combine sensory evidence with prior expectations in a precision-weighted manner. Although Bayesian inference provides a clear normative account of how such integration should occur, the neural mechanisms through which the brain represents and combines priors and likelihoods remain poorly understood. Across two preregistered experiments, we investigated how the precision of prior expectations and sensory likelihoods influences visual motion judgements and associated neural activity patterns during a random-dot motion estimation task. Neurotypical adult participants (N=80, 58 female) reported directions of visual motion stimuli, with motion coherence varying randomly across trials. Prior expectations were manipulated block-wise by varying the probabilities with which different motion directions were presented. Consistent with precision-weighted inference, response accuracy improved as coherence increased with robust response biases toward the expected motion direction. Neural activity measured using electroencephalography (EEG) revealed reliable effects of prior expectation on the univariate central-parietal positivity (CPP), consistent with reduced accumulation of sensory evidence under informative priors. Multivariate analysis using inverted-encoding models revealed robust effects of prior informativeness on motion-specific neural representations, but only late, during response planning stages. Together, these findings demonstrate that precision-weighted inference primarily occurs at late stages of the decision process and challenge predictive-processing accounts that emphasise early sensory processing.

**Significance Statement:** Perceptual decisions require combining uncertain sensory information with prior expectations about what is likely to occur. Although behaviour often follows this principle, it remains unclear how the brain integrates prior expectations with incoming sensory evidence. Using a visual motion task and concurrent brain imaging, we show that expectations do not alter early sensory processing but instead influence later stages of decision formation and action planning. Neural representations of sensory evidence primarily reflect stimulus reliability, whereas prior expectations selectively enhance representations during response preparation. These findings challenge influential theories proposing that expectations affect early sensory processing and instead highlight their contribution to later decision-related processes. This work advances understanding of how the brain uses experience to optimise perceptual decisions under uncertainty.

## Introduction

Adaptive interaction with the environment requires making perceptual decisions in the face of uncertainty. The Bayesian inference framework offers a principled account of how such decisions are mediated; the brain combines sensory evidence with prior expectations to infer the most likely state of the world (Gold & Stocker, 2017; Körding & Wolpert, 2004; Ma et al., 2023; Weiss et al., 2002). Importantly, this combination is not a simple average. Instead, the contribution of each source of information is weighted by its precision. Despite its success in describing behaviour, however, it remains unclear how precision-weighted integration is achieved in the brain. We investigated this question across two preregistered experiments where adult human participants learned to use priors to guide their choices while we recorded their brain activity using electroencephalography (EEG).

Under accumulate-to-bound models such as drift-diffusion (Ratcliff & McKoon, 2008), priors for choice alternatives are integrated into the decision process as response bias in favour of the more likely alternative (Mulder et al., 2012; Summerfield & de Lange, 2014). Neural evidence for this bias has been identified in increased preparatory motor activity in both humans and non-human animals (Basso & Wurtz, 1997; De Lange et al., 2013; Kelly et al., 2021; Rao et al., 2012; Walsh et al., 2024). However, recent accounts suggest that priors also increase the *rate* at which evidence is accumulated by directly impacting representations of expected stimuli (Diaz et al., 2024; Kelly et al., 2021; Walsh et al., 2024). Under this view, the brain utilises prior expectations by proactively modulating neural populations tuned to the expected stimulus values (de Lange et al., 2018; Press et al., 2020; Summerfield & de Lange, 2014). Examinations of population-level neural activity patterns show that stimulus representations can be biased toward expected values (Kok et al., 2013; Park et al., 2023) and are increasingly precise the closer they are to the expected value (Kok et al., 2012; Yon et al., 2018). Moreover, early neural responses associated with an expected stimulus can occur when no stimulus is presented (Kok et al., 2014; Todorovic & de Lange, 2012), and may even be activated *before* a stimulus appears (Aitken et al., 2020; Blom et al., 2020; Feuerriegel, Blom, et al., 2021; Kok et al., 2017). These findings point to an early modulation of sensory representations by priors, consistent with influential predictive processing theories (de Lange et al., 2018; Press et al., 2020; Summerfield & de Lange, 2014; Walsh et al., 2020).

Despite these advances, it remains unclear whether the influence of prior expectation on neural stimulus representations scales with the precision of that prior. Moreover, it remains unknown whether such effects interact with sensory precision, thereby approximating Bayesian inference at the level of neural representation. We addressed these questions in two preregistered experiments where participants learned and used priors to guide choices in a random-dot motion estimation task. Using a factorial design we parametrically manipulated the precision of prior expectations and sensory signals, and evaluated their effects on behavioural error distributions. Capitalising on the high temporal resolution of EEG combined with multivariate population tuning analyses, we characterised how and when neural representations of motion direction are impacted by prior and likelihood precision. If, as suggested by predictive processing models (de Lange et al., 2018; Press et al., 2020; Walsh et al., 2020), priors are integrated into the decision process by either sharpening or biasing the encoding of decision-relevant stimuli, then increasing the precision of priors should increase the accuracy and/or bias of neural representations decoded from EEG. This effect should arise at early timepoints around stimulus onset and be at least as strong as expectation effects on representations later in the trial. Finally, if these processes underlie the approximation of Bayesian inference regularly observed in behavioural studies (Chalk et al., 2010; Körding & Wolpert, 2004), then expectations should interact with stimulus precision such that they exert the strongest influence when the stimulus is least precise.

## Materials & Methods

### Participants

Forty-three volunteers participated in the behavioural experiment (Experiment 1). Two did not finish the task, and one was excluded for poor task performance (see *Behavioural Analyses*) resulting in a final sample size of 40 participants for Experiment 1 (33 female, 37 right-handed, 18 – 38 years). Another 43 volunteers participated in the EEG experiment (Experiment 2). One volunteer withdrew after the first EEG session and two were excluded for poor task performance, leaving a final sample of 40 participants in Experiment 2 (25 female, 38 right-handed, 18 – 37 years). All participants self-reported normal or corrected-to-normal vision, provided written informed consent and were compensated $20 AUD per hour for their time. The sample size of N = 40 per experiment is consistent with previous work from our group that used similar tasks and analysis techniques (Rangelov et al., 2021, 2024; Rangelov & Mattingley, 2020). The study was approved by The University of Queensland Human Research Ethics Committee (HE001494). Both experiments were pre-registered with the Open Science Framework (Experiment 1: osf.io/s7d59; Experiment 2: osf.io/pt82w).

### Stimuli, task & procedure

Noisy motion signals were generated using a random dot-motion stimulus. The stimulus comprised 200 white dots (0.05 degrees radius) pseudo-randomly distributed around a yellow fixation point (0.05 degrees radius) within a circular aperture (8 degrees radius) against a black background. Dot positions were updated every frame to generate apparent motion. The direction of motion for each dot was sampled randomly and independently from a von Mises distribution (circular Gaussian) defined by a mean (µ_s_) and a precision parameter (κ_s_). The precision of motion information available in the stimulus was manipulated by varying κ_s,_ between high (0.4) and low (0.2) in Experiment 1 (**Figure 1B**). Dot speed was held constant at 3 degrees visual angle (dva) per second. Dots had an infinite lifetime and were wrapped around if they left the stimulus aperture.

**Figure 1.**
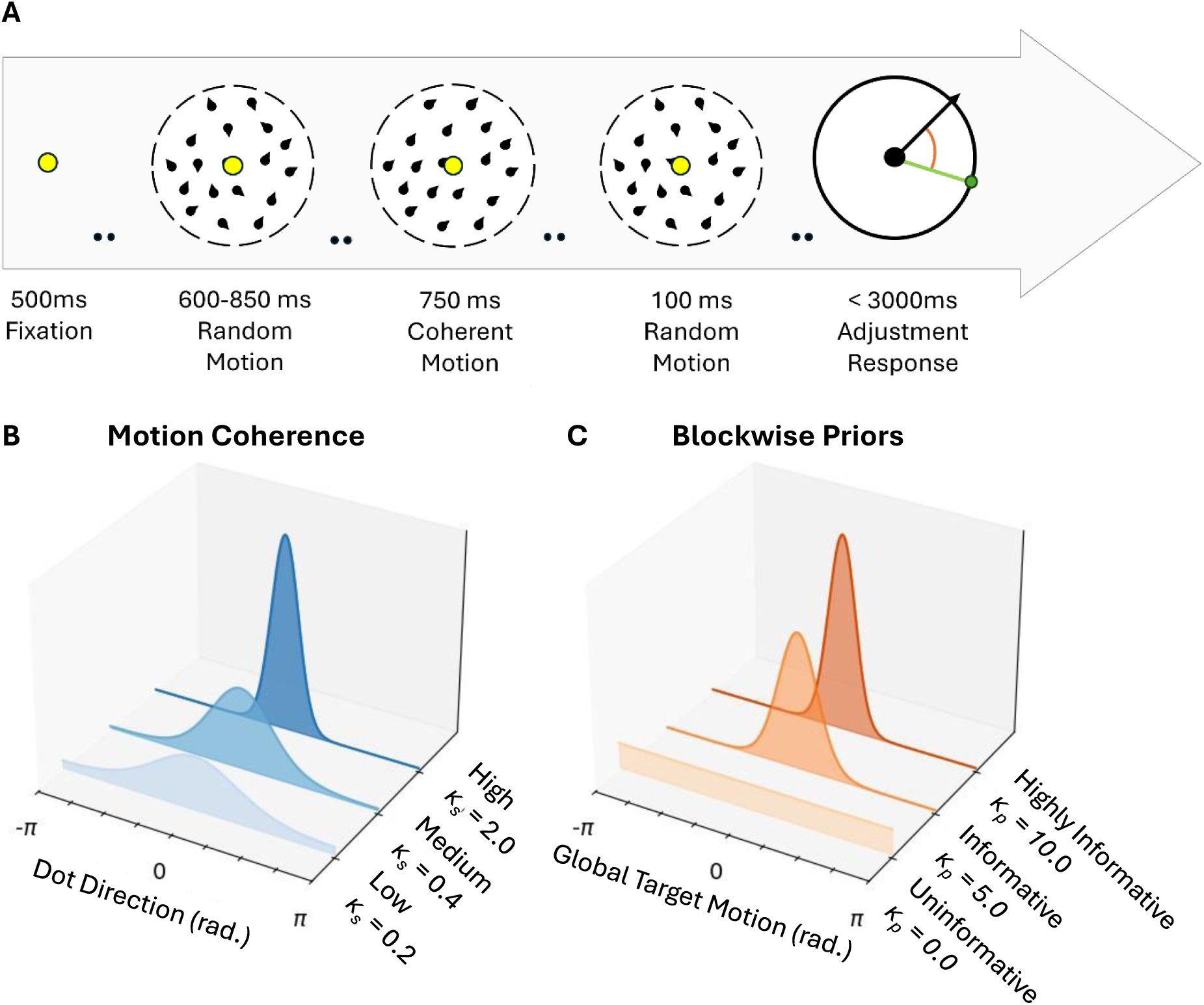
Overview of the task and the manipulation of stimulus and prior precision. **A)** Structure of a single trial from pre-fixation to response. Note: trial diagram is schematic only, colours and scale differ from actual design (see Methods). **B)** Stimulus precision was manipulated via motion coherence, corresponding to the precision of the sampling distribution of possible dot-directions on each frame. Zero corresponds to the global target motion direction. **C)** Priors for motion direction were manipulated via the block-level sampling distribution for global target motion directions. These distributions could either be completely uninformative (uniform) or contain some information about the probability of upcoming stimulus values. In this case zero corresponds to the central tendency of the block-wise sampling distribution/prior.

To manipulate priors, the global motion signal (i.e., the target) on each trial was sampled from a block-level hyper-distribution, which was also a von Mises distribution but defined by a different mean (µ_p_) and precision (κ_p_; **Figure 1C**). Each participant learned priors for three motion directions (µ_p_) separated by 120 degrees. Priors were either uninformative (κ_p_ = 0, effectively uniform), weakly informative (κ_p_ = 5) or strongly informative (κ_p_ = 10). These hyperparameters varied across blocks and counterbalanced such that participants experienced every combination of µ_p_ and κ_p_ twice, yet no two consecutive blocks shared a value of µ_p_. Between participants, µ_p_ angles were adjusted by a small amount to ensure even sampling around the full circle at the group level.

Each participant completed 18 blocks of 60 trials. Each trial (see **Figure 1A**) began with a fixation point (500 ms), followed by a dot-display containing only random motion (κ_s_ = 0). After a variable period of 600-850 ms, the target motion direction was presented for 750 ms before motion returned to random for a final 100 ms. Participants were instructed to estimate the target motion direction within 3000 ms using an on-screen response dial presented at fixation. The dial comprised a black circle with a red outline (4 degrees radius) and a yellow ‘arm’ allowing 360 degrees of movement around the circle. The initial angle of the arm on each trial was randomly sampled from a uniform distribution of values ranging from –π to π radians so participants could not start planning a response before the onset of the response display.

The main aim of Experiment 2 was to characterize the neural correlates of sensory processing and decision-making using both univariate and multivariate analyses of brain activity measured using EEG. Aside from the minor departures mentioned below, the protocol for Experiment 2 was identical to that of Experiment 1. To accommodate the extra statistical power required for multivariate EEG analysis, participants in Experiment 2 completed 2160 trials (36 blocks) spread over two sessions. Since participants had more exposure to the prior statistics in Experiment 2, we rotated the values of µ_p_ 60 by degrees between sessions 1 and 2. In addition, we introduced a ‘high’ motion coherence condition (κ_s_ = 2.0) which introduced strong motion-selective neural responses that were subsequently used for training the EEG decoding models. Experiment 2 therefore had three levels of motion coherence (0.2, 0.4 and 2.0; “low”, “medium” and “high” coherence, respectively). Since we planned to train decoding models on high coherence trials from the uninformative prior condition, we wanted to avoid the possibility that these models might become biased because of spurious failures of randomization during sampling. To eliminate this possibility, we changed the way motion directions were selected in the uninformative-prior condition (κ_p_ = 0) such that target directions were now drawn without replacement from a pre-determined set of 60 evenly spaced directions spanning the full circle.

### Apparatus

The experiments were conducted in a dark room using a Dell Precision T1700 computer with either a 32” Display++ LCD monitor (Cambridge Research Systems Ltd., Rochester, UK) (Experiment 1) or a 24” ASUS VG248 (ASUSTeK Computer Inc., Taiwan) (Experiment 2). Both displays had a 1920*1080-pixel resolution and a 120 Hz refresh rate. Stimuli were generated using custom Python code (Python version 3.8.10) using the PsychoPy toolbox (v2021.1.3; Peirce et al., 2019). EEG signals were recorded using 64 Ag-AgCl electrodes (ActiveThree, BioSemi) arranged in the 10-20 layout and sampled at 1024 Hz.

### Behavioural Analyses

To remove any influence of pre-emptive responding, we removed trials with response times shorter than 200 ms. Trials where the participant did not response within 3000 ms were also removed. Per our preregistered criteria, two participants for whom more than 20% of the total trial count were removed for the above reasons were excluded from Experiment 2. Fewer than 0.1% of trials were removed from the remaining sample in either experiment. We also set a generous lower bound on acceptable task performance; participants were only excluded from analysis if the circular standard deviation of response errors was greater than π / 2 radians. One participant was excluded from Experiment 1 for this reason.

We characterized behavioural performance in the estimation task by examining histograms of response errors per participant across experimental conditions. Errors were calculated via circular subtraction of veridical motion directions from responses. Response accuracy for each trial was quantified using the absolute magnitude of errors subtracted from one. Bias toward a learned prior was quantified by re-coding the sign (+/-) of errors such that those made toward µ_p_ became positive, and those away became negative. We term this operationalisation of bias the *signed error.* Attractive bias toward µ_p_ is thus captured by a positive shift in the circular mean of the distribution of signed errors.

### EEG Analyses

EEG signals were processed in Python (version 3.12.7) using the MNE analysis toolbox (version 1.8.0; Larson et al., 2024). The data were referenced offline to the average electrode, band-pass filtered between 0.2 Hz and 40 Hz and notch filtered at 50 Hz to eliminate line noise. Signals were then epoched between –0.95 s and 2.0 s relative to onset of the motion signal, baselined relative to the average signal during a 100ms period of random motion (–0.35 s to –0.25 s) prior to target onset and down-sampled to 256 Hz. The baseline period was selected to minimize the possibility of artifacts introduced by anticipation of target motion-onset which may have resulted from the variable duration of pre-target random motion. The epoched data were next submitted to an automated preprocessing pipeline for artefact detection and correction using the FASTER algorithm (Nolan et al., 2010). Between 26 and 100 trials per participant were identified as EEG outliers and removed from further analyses (median = 61 trials).

Decision-relevant accumulate-to-bound dynamics have previously been observed in event-related build-ups in positive voltage over central and parietal cortices (O’Connell et al., 2012). To analyse these centro-parietal positivity (CPP) dynamics, we extracted motion-locked event-related potentials by averaging over a cluster of 5 electrodes at the center of the head (Cz, CPz, Pz, CP1 & CP2) – aligning with scalp positions reported in previous studies of perceptual decision-making (O’Connell et al., 2012; Rangelov & Mattingley, 2020; Stone et al., 2024).

We used two distinct computational approaches to model feature-specific information about motion direction in terms of participants’ EEG activity. First, we characterized the sensitivity of individual electrodes to different motion directions by estimating the linear relationship between motion direction and electrode voltage. Using Equation 1, we regressed the voltage *V* from electrode *i* and time sample *t* against the sine and cosine transform of the presented motion direction *θ* using a general linear mixed-effects model which allowed random intercepts *u* per participant ID *j*.

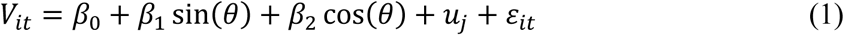

We then indexed motion sensitivity *S* of electrode *i* at time *t* as the vector length defined by the fitted *β* values for each term (Equation 2).

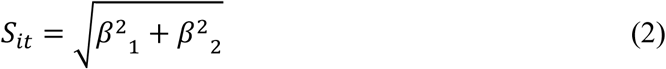

Since *S* was always positive, we permuted this analysis 1,000 times with shuffled values of *θ* to generate an empirical null distribution from which we could estimate a p-value for every electrode and timepoint. We restricted this analysis to high motion coherence trials from blocks where the prior was uniform to ensure comparability with the multivariate inverted-encoding analysis, as described below.

In a second analysis, we employed an inverted-encoding model to characterize multivariate patterns of feature-specific neural responses across the entire electrode population (Brouwer & Heeger, 2009; Kok et al., 2017; Rangelov et al., 2021). Briefly, this technique uses ordinary least squares to map multivariate EEG sensor activity to hypothetical motion-tuned channel responses (i.e., estimating a forward model), then inverts this mapping to recover population-tuning profiles from held-out data.

Consistent with previous work using motion stimuli (Rangelov et al., 2021), the forward model comprised 16 hypothetical motion-responsive channels, with evenly distributed tuning preferences between –π and π radians. Each channel consisted of a half-wave rectified sinusoid raised to the fifth power. Hypothesized motion tuning for each target direction could then be expressed as the weighted circular sum of the 16 channels. EEG activity for each stimulus presentation could thus be described using the linear model expressed in Equation 3:

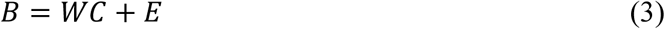

Where *B* represents the EEG data as a *m* sensors x *n* trials matrix, *W* is a *m* sensors x 16 motion channel weight matrix that describes the transformation from EEG activity to motion direction, *C* indicates the hypothesized channel activities (16 motion channels x *n* trials), and *E* is the residual term.

To compute the inverse model, which takes EEG sensor activity back to motion channel activations, we estimated weights that reproduced the hypothesized channel activities with the least error via ordinary least squares optimization. Using a procedure described in previous work (Aitken et al., 2020; Kok et al., 2017; Rideaux et al., 2023), we accounted for highly correlated EEG sensor activity by incorporating noise-covariance into the weight estimation procedure, regularized using an analytically determined optimal shrinkage parameter (Ledoit & Wolf, 2004).

We conducted this analysis across trials for each participant and time sample. However, because the structure of our experimental task involved non-uniform sampling of motion directions, we estimated the inverse models only on trials sampled from a uniform distribution and presented with high motion coherence. We then used these models to recover motion tuning in the other conditions. Selectively splitting trials into these training and testing sets meant that tuning properties recovered from held-out data only reflect properties of the neural stimulus representations, rather than factors arising from non-uniform sampling. Since these training trials also served as an important control condition in our inferential analyses, we additionally estimated their motion tuning using a leave-one-out cross validation procedure.

To decode the stimulus direction represented in EEG activity on each trial, we used an aggregate index of the channel response profile by computing the dot product of the 16 recovered channel activities with their respective complex-valued motion tuning preferences. The angle of the resulting vector reflects the decoded motion direction, and the vector length provides an estimate of the single trial tuning width. We then quantified single-trial accuracy as the cosine of the neural error magnitude, expressed as the difference between the decoded direction and the stimulus direction (the same as behavioural errors). An accuracy score of 1 indicates perfect decoding and a score of 0 indicates chance. Next, we calculated neural bias by applying the same sign-flipping procedure previously described for behavioural response errors to the recovered neural errors.

We also examined whether stimulus representations were stable across time, and whether the representations that were active during encoding of coherent motion might also extend to time periods either before or after the period where coherent motion information was physically presented to the participant. These analyses involved a temporal generalization approach whereby inverse models obtained at each timepoint from the training set were applied at every other timepoint during the trial. This approach permitted us to create two-dimensional matrices of decoding accuracy for each experimental condition (King & Dehaene, 2014; Kok et al., 2017; Mostert et al., 2015).

### Experimental Design & Statistical Analyses

The main independent variables were stimulus precision (κ_s_ = 0.2, 0.4 and 2.0) and prior precision (κ_p_ = 0, 5 and 10), resulting in 2×3 and 3×3 within-subject designs for Experiments 1 and 2, respectively. Statistical analyses for behavioural accuracy and bias were performed at the single-trial level using mixed-effects general linear models implemented with the *statsmodels* package in Python (*statsmodels* version 0.14.4; Seabold & Perktold, 2010). Each model included fixed-effect predictors of stimulus precision (*κ*_s_) and prior precision (*κ*_p_) and their interaction, while allowing random intercepts across participants:

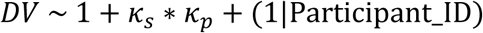

Time-resolved statistical analyses of neural data (CPP amplitude, decoding accuracy and bias) were similarly performed at the single-trial level using the same approach repeated for each timepoint. In this case the estimated parameters were assessed for statistical significance against 0 using the false discovery rate to control for inflation of false-positive probability due to multiple comparisons (Benjamini & Hochberg, 1995). Post-hoc contrasts between experimental conditions involved one-sample t-tests of difference scores against zero, corrected for multiple comparisons using cluster-based permutation-tests (*p* < .01 cluster admission threshold; clusters with *p* < .05 after 1000 permutations are reported).

## Results

### Motion Perception Reflects a Precision-Weighted Inference Strategy

Response errors in Experiment 1 were unimodal and clustered around zero, indicating good task performance at each level of stimulus precision (**Figure 2A**). As expected, response accuracy increased in step with stimulus precision (main effect of precision: *β* = 1.37, *SE* = 0.038, *t*(42898) = 36.25, *p* < 0.001, **Figure 2B**).

**Figure 2.**
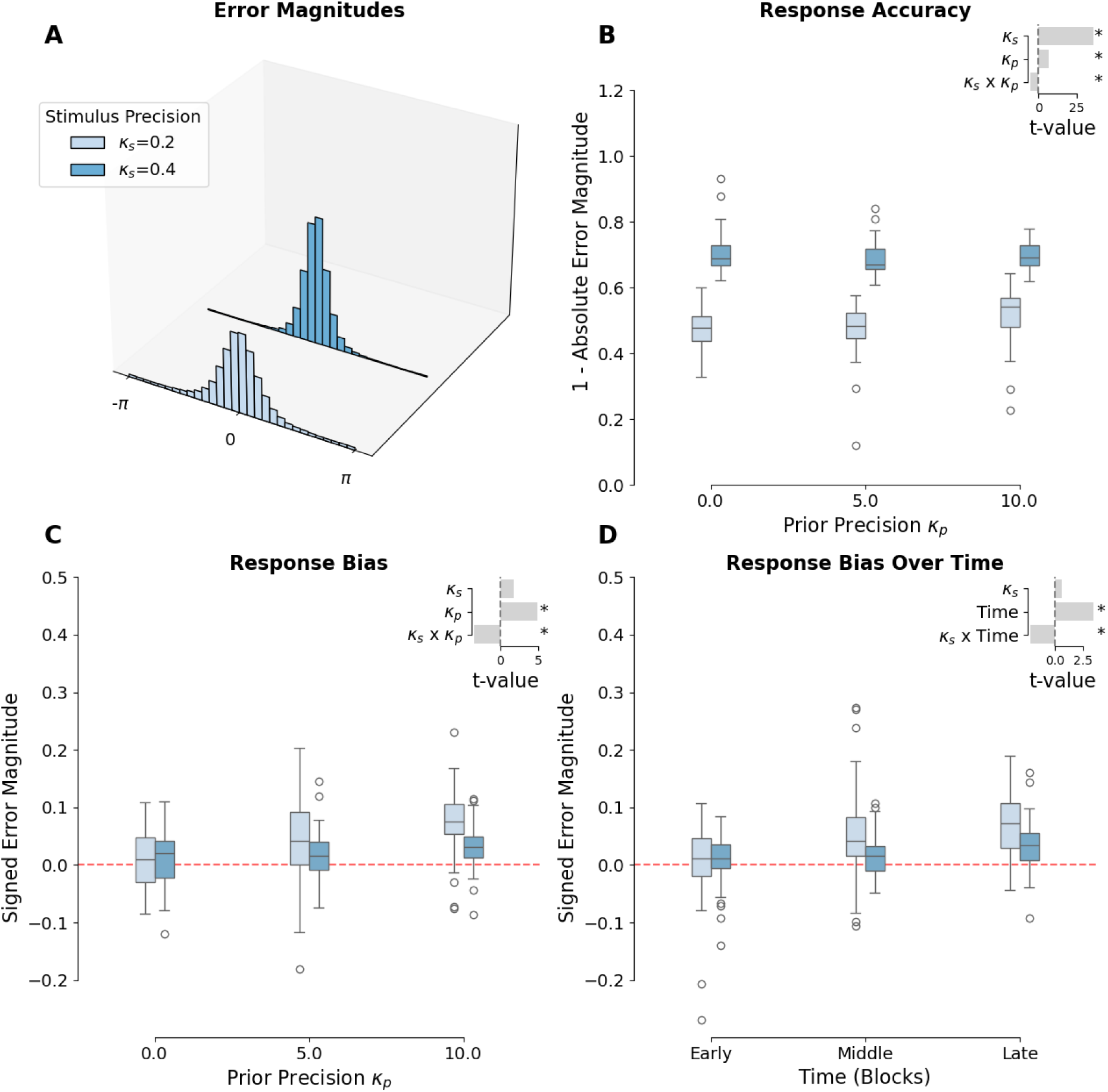
Results of Experiment 1 (Behavioural experiment). Trial-level error histograms for each level of stimulus precision. **B)** Response accuracy stratified by stimulus precision and prior precision. **C)** Response bias stratified by stimulus precision and prior precision. Bias expressed as mean signed-error magnitudes such that attraction to the central-tendency of the current prior is captured by a positive shift in the error distribution. **D)** Bias stratified by stimulus precision and time within the session grouped into three equal-sized bins. Here, time reflects the exposure participants had to each of the priors. For visualisation, the data used to construct boxplots were normalised to remove the between subjects’ variability (Cousineau, 2005). Inset bar plots indicate t-statistics and significance of mixed GLM parameter estimates. * = p < .05.

There was also a comparatively small, but statistically significant, main effect of prior precision such that stronger priors resulted in more accurate responses (*β* = 0.012, *SE* = 0.002, *t*(42898) = 6.42, *p* < 0.001). Importantly, the interaction between stimulus and prior precision was significant (*β* = –0.03, *SE* = 0.006, *t*(42898) = –5.19, *p* < 0.001), such that the improvement in accuracy afforded by a stronger prior was more pronounced for the least precise coherent motion stimuli.

Response bias analyses revealed attractive biases toward expected motion directions (i.e., positive shifts) with stronger priors yielding stronger attractive biases (*β* = 0.013, *SE* = 0.003, *t*(42898) = 4.93, *p* < 0.001, **Figure 2C**). There was no main effect of stimulus precision on response bias. Consistent with our preregistered hypotheses, the effect of prior precision significantly interacted with stimulus precision (*β* = –0.029, *SE* = 0.008, *t*(42898) = –3.44, *p* < 0.01) such that attraction toward the prior was maximal when the stimulus was weakest and the prior strongest.

Next, we asked how priors were learned and maintained over the course of a behavioural session. Though the prior in each block was always different to that of the previous block, three central tendencies (i.e., prior mean; µ_p_) were repeated throughout each session. We might expect, therefore, that response biases should be stronger in later, relative to earlier blocks given participants’ increased exposure to the prior. To explore this question, we divided the data into three sets depending on whether they were early, in the middle or late within each experimental session, and then modelled bias as a function of stimulus precision and time bin (i.e., prior exposure). This analysis revealed that biases did indeed increase significantly with greater exposure to the prior (*β* = 0.045, *SE* = 0.013, *t*(42898) = 3.45, *p* < 0.01; **Figure 2D**). Exposure to the prior also significantly interacted with stimulus precision in both experiments such that the strongest biases emerged when stimulus precision was low (*β* = –0.090, *SE* = 0.042, *t*(42898) = –2.15, *p* < 0.05), suggesting participants weighted priors more heavily after prolonged exposure.

Experiment 2 served as a replication of the behavioural results of Experiment 1, but now with neural activity recorded concurrently via EEG. We replicated each of the key behavioural effects in Experiment 2 (**Figure 3**): response accuracy was significantly predicted by both stimulus (*β* = 0.199, *SE* = 0.003, *t*(85521) = 67.16, *p* < .001) and prior precision (*β* = 0.006, *SE* = 0.001, *t*(85521) = 11.6, *p* < 0.001) and by their interaction (*β* = – 0.003, *SE* < 0.001, *t*(85521) = –6.75, *p* < 0.001; **Figure 3B**) such that responses to the weakest stimuli were most improved under stronger priors. Similarly, the magnitude of response biases were predicted by the precision of the learned prior (*β* = 0.003, *SE* = 0.001, *t*(85521) = 4.73, *p* < 0.001) and its interaction with stimulus precision (*β* = –0.003, *SE* = 0.001, *t*(85521) = –4.37, *p* < 0.001; **Figure 3C**). Once again, we found that blocks later in each experimental session exhibited greater biases relative to those toward the beginning (*β* = 0.023, *SE* = 0.004, *t*(85521) = 5.80, *p* < 0.001; **Figure 3D**) indicating priors played an increased role in decision formation as they were learned over time. Exposure to the task statistics also interacted with stimulus precision (*β* = –0.008, *SE* = 0.003, *t*(85521) = –2.44, *p* < 0.05) suggesting priors were weighted more heavily as familiarity with them increased and as stimulus precision decreased.

**Figure 3.**
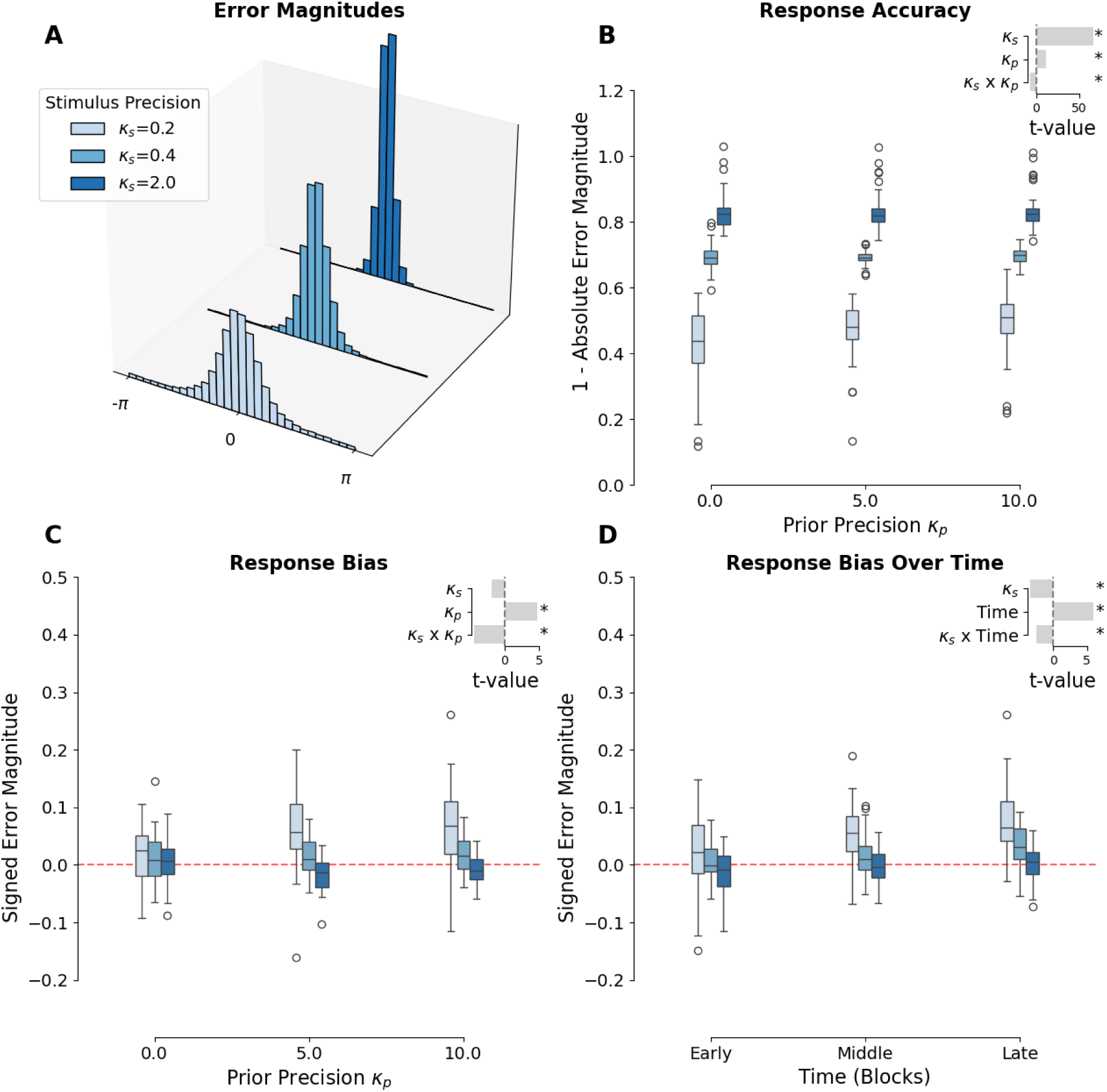
Behavioural results for Experiment 2 (EEG + Behaviour). **A)** Trial-level error histograms for each level of stimulus precision. **B)** Response accuracy stratified by stimulus precision and prior precision. **C)** Response bias stratified by stimulus precision and prior precision. Bias expressed as mean signed-error magnitudes such that attraction to the central-tendency of the current prior is captured by a positive shift in the error distribution. **D)** Bias stratified by stimulus precision and time within the session grouped into three equal-sized bins. Here, time reflects the exposure participants had to each of the priors. For visualisation, the data used to construct boxplots were normalised to remove the between subjects’ variability (Cousineau, 2005). Inset bar plots indicate t-statistics and significance of mixed GLM parameter estimates. * = p < .05.

Taken together, the behavioural results of Experiments 1 and 2 are consistent with a precision-weighted Bayesian inference strategy whereby observers optimize their decision-making by integrating probabilistic representations of the sensory input with expectations about the task. Importantly, the robust interactions found between precision of motion signals and informativeness of priors strongly suggest that observers utilize the precision of their prior expectations, as well as that of the available sensory evidence, when making decisions about motion direction.

### Priors and Likelihoods Differentially Influence Neural Signatures of Evidence Accumulation

Having established that participants’ decision-making behaviour was sensitive to both the reliability of sensory evidence and learned priors, we next examined the associated patterns of brain activity, as reflected in the central parietal positivity (CPP) component of participants’ EEG signals, a well-studied neural correlate of decision-making. Previous work has established that the CPP tracks accumulated sensory evidence relevant to an upcoming decision regardless of the sensory modality involved (O’Connell et al., 2012). In our task, the CPP followed the expected time course of evidence accumulation, featuring a ramping of neural activity ∼ 200 ms after the onset of coherent motion which peaked between 300 and 400 ms depending on the strength of sensory evidence (**Figure 4A**). These observations were supported by time-resolved linear modelling which revealed a strong effect of stimulus precision such that CPP amplitudes increased markedly in step with stimulus precision (**Figure 4C**). This result is in line with previous work showing that the CCP is sensitive to the amount of evidence available in the target for decision-making (Kelly & O’Connell, 2013; Rangelov et al., 2021; Stone et al., 2024), and suggests that the rate of evidence accumulation in the visual motion task was facilitated by stimulus precision.

**Figure 4.**
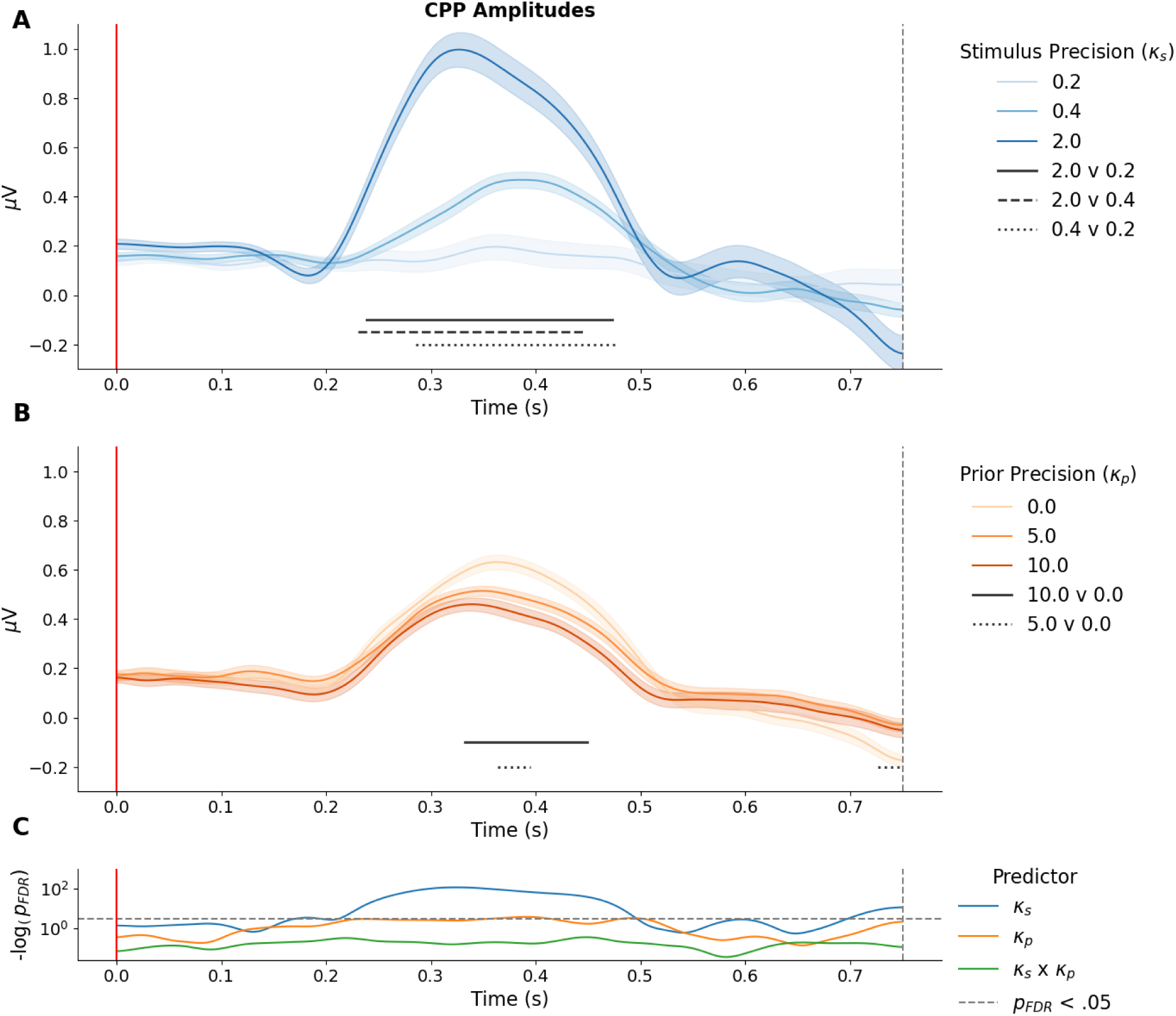
Mean motion-locked central parietal positivity (CPP) recorded during the 750 ms period of coherent motion. Plots are stratified by **A)** motion coherence and **B)** prior precision. Shaded regions reflect the within-participant standard error of the mean. Vertical red lines and grey dashed lines indicate coherent motion onset, and offset, respectively. Black solid, dashed and dotted lines indicate cluster significance p < .05 of pairwise condition comparisons. **C)** Log-transformed p values for fixed effects of stimulus and prior precisions and their interaction on CPP amplitude, presented on a log scale. Horizontal dashes indicate the log-transformed significance threshold. For the purposes of presentation, each trace was smoothed using a temporal Gaussian filter (sigma = 4). All statistical comparisons were performed on the unsmoothed data.

We also observed a small and transient, but significant effect of prior precision on CPP amplitude, whereby neural responses under informative priors (κ_p_ = 5 & κ_p_ = 10) were reduced relative to the uniform prior condition (κ_p_ = 0; **Figure 4B**). This finding suggests that the brain accumulates less evidence when stimulus values are predictable compared with when they are random, consistent with the accumulate-to-bound decision-making framework.

Here, a non-uniform prior over potential decision outcomes shifts the starting point of evidence accumulation in favor of more likely target values. This shift ultimately results in an imbalance in the amount of evidence accumulation necessary to trigger the favored decision. The slope of the CPP is frequently found to be proportionate to the quality of sensory evidence, i.e., the stimulus representation (Kelly & O’Connell, 2013; Rangelov et al., 2021; Stone et al., 2024), an effect clearly reflected in the gradation of slopes when stratifying CPP waveforms by motion coherence (**Figure 4A**). However, this same pattern was not apparent when splitting the CPP by the strength of the learned prior (**Figure 4B**), suggesting little or no influence of a prior on the quality of evidence being sampled.

### Feature Specific Decoding Analyses

To test whether prior expectations affected the neural representation of motion direction during the task, we undertook two analyses that quantified the spatiotemporal dynamics and representational fidelity of motion-specific stimulus information. Time-resolved linear modelling of single-electrode activities revealed a posterior-anterior progression of motion-selective information across the scalp as participants observed and then acted upon the motion stimulus (**Figure 5A**). Motion-direction information was initially present over central and posterior areas (∼400 ms post stimulus) before progressing into central and frontal areas toward the end of the observation period (∼700 ms). Early during the response period (∼1000 ms), motion-direction information was evident over nearly all electrode sites including over posterior areas, consistent with maintenance (or re-activation) of sensory representations during decision-formation and response. Late activity in the lead-up to the response (average response time ∼ 1800 ms) was bilaterally localised over frontal electrode sites. Importantly, while participants were instructed to maintain fixation and keep their response hand still during stimulus observation, they were free to make saccades and move the computer mouse during the response period. It is thus possible these later signals reflect stimulus-correlated motor-activity as well as stimulus-specific neural activity.

**Figure 5.**
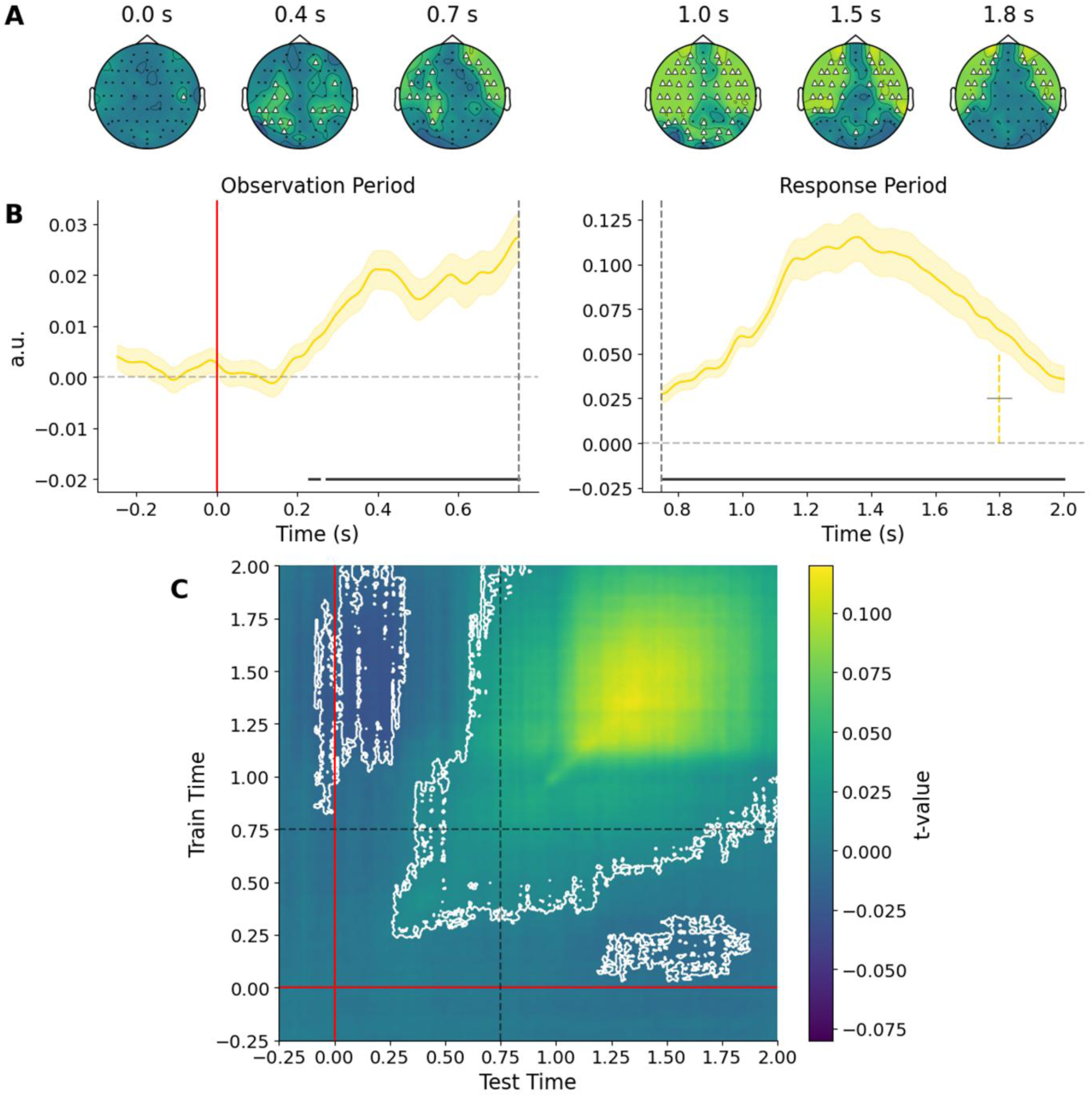
Motion-specific neural responses to coherent motion signals. **A)** Individual electrode sensitivity to motion information. Here, motion sensitivity is expressed as negative log transformed p-values, corrected for multiple comparisons using the false discovery rate over both time and electrode dimensions. Electrode sites highlighted by triangles indicate p_FDR_ < .05. **B)** Time course of neural tuning to motion direction derived from the inverted-encoding analysis. Motion tuning is expressed as average decoding accuracy during observation of coherent motion (left panel) and response (right panel; note the different y-axis scale). Shaded regions indicate the group-level standard error, and the black line denotes clusters where accuracy is significantly above chance level (horizontal grey dashes). Coloured vertical bar (dashed) in the response panel corresponds to the mean group-level response time with the grey bar denoting standard error. **C)** Temporal generalisation decoding matrix averaged over all condition combinations. Pixel values correspond to t-statistics describing difference between decoding accuracy and chance. White contours denote cluster significance following 10000 permutations. As above, Gaussian smoothing was applied to time series for plotting only.

In a second analysis, we trained inverted encoding models to characterise the dynamics of motion-tuning throughout the decision-process from whole-scalp EEG activity. We expressed the strength of neural representations within each condition as the aggregated accuracy of the decoded motion directions on each trial. As depicted in **Figure 5B**, stimulus motion was robustly decoded from approximately 200 ms post stimulus onset through to (and beyond) the average response time (∼ 1800ms). Temporal generalisation analysis confirmed that after an initial ramping-up of motion-specific activity, the representation encoded during observation of coherent motion was maintained well into the response period (see **Figure 5C**; persistent decoding cluster including off diagonal extension from testing at ∼300 ms through to ∼ 1200 ms).

Next, we asked whether neural tuning to motion direction at each timepoint was affected by our manipulations of stimulus and prior precision. Trial-level linear mixed-effects modelling and pairwise comparison of the decoding accuracy time series revealed robust differences in the precision of neural representations as a function of stimulus precision during both observation of the coherent motion target and response generation (**Figure 6A**). The differences during motion observation mirror the order observed in the univariate CPP data (**Figure 4A**), corroborating our interpretation that the quality of evidence accumulated is proportionate to the robustness of participants’ representation of the stimulus. Importantly, motion tuning during this motion observation period was not modulated by the precision of the prior (**Figure 6B**), reinforcing the interpretation that recently learned priors impact the starting point of the decision process, rather than directly impacting the encoding of sensory evidence. In contrast, prior precision did significantly influence motion tuning during the response period; decoding accuracy from ∼1000-1700 ms increased in step with the reliability of the current prior, suggesting that representations of stimuli are enhanced according to their prior probability during decision-making and action preparation.

**Figure 6.**
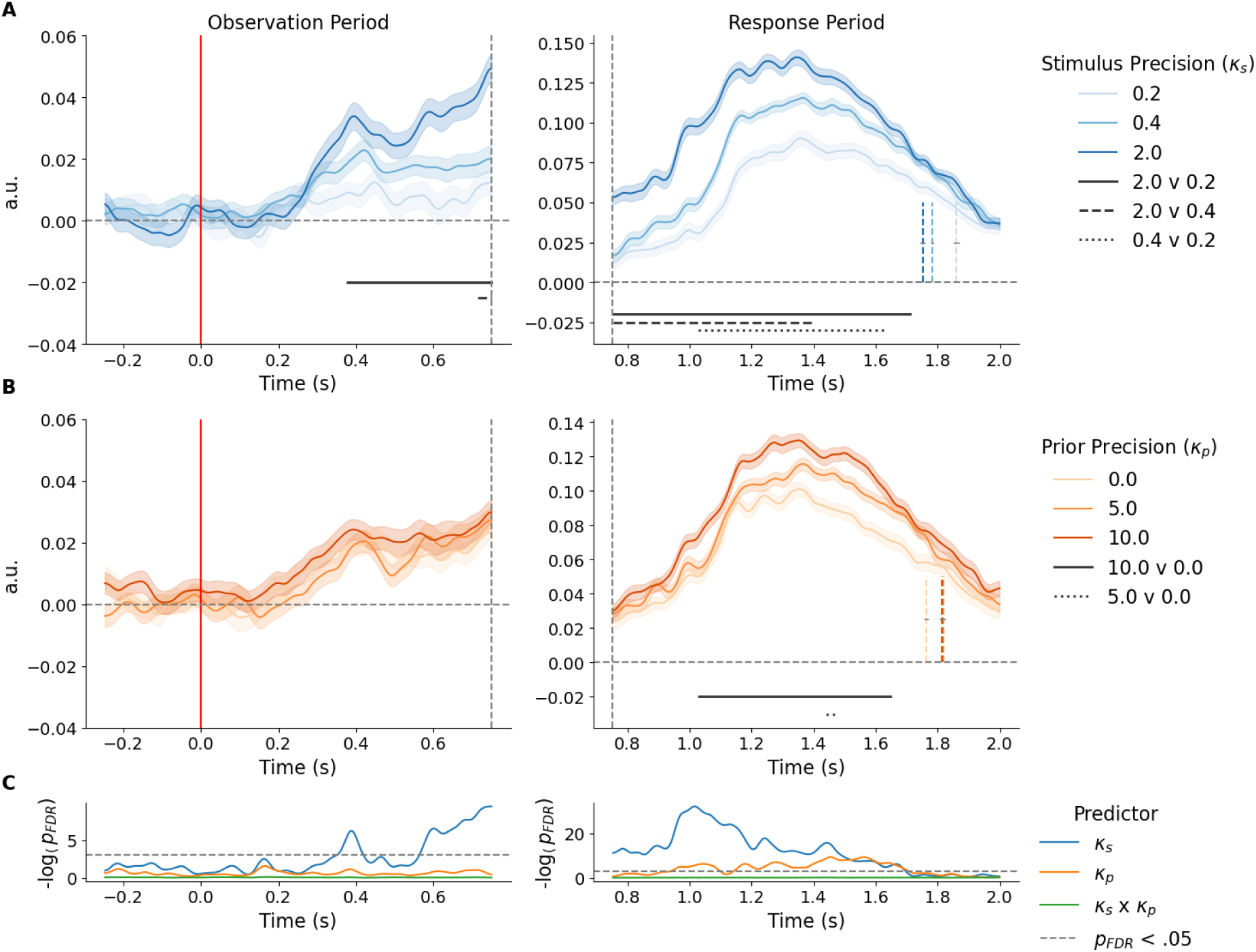
Neural tuning to motion direction expressed as decoding accuracy stratified by **A)** stimulus precision and **B)** prior precision during the observation and response periods (note the difference y-axis scales). Vertical red lines and grey dashed lines indicate coherent motion onset, and offset, respectively. Black solid, dashed and dotted lines indicate cluster significance p < .05 of pairwise condition comparisons. **C)** Log-transformed p values for fixed effects of stimulus and prior precisions and their interaction on decoding accuracy. Horizontal dashes indicate the log-transformed significance threshold. For the purposes of presentation, each trace was smoothed using a temporal Gaussian filter (sigma = 4). All statistical comparisons were performed on the unsmoothed data.

Bayesian decision-theory dictates that the best strategy for minimising errors is to make estimates that are biased toward the central tendency of the prior (with the degree of bias weighted by the respective precisions of the prior and sensory likelihood (Körding & Wolpert, 2004; Ma et al., 2023)). Since we observed such biases in the behavioural data (**Figure 3C**), we might expect that the largest impact of a strong prior should be to shift the value of the represented motion direction in its favour. To investigate this possibility, we quantified neural bias at the trial-level using the same method as for behavioural bias by recoding the error sign to reflect prior attraction. No systematic shifts in the represented motion direction were observed with respect to the prior in either the observation or response periods suggesting that, on average, motion representations faithfully tracked the presented target direction even once the stimulus was no longer visible (see **Figure S1**).

The analyses reported so far reveal robust differences in the fidelity of motion-specific patterns of brain activity as a function of the precision of both stimuli and learned priors during response preparation and execution (**Figure 6**). While the effect of stimulus precision is already apparent during observation of the coherent motion target, the effect of prior precision only emerged later during the response period. To respond, participants rotated an on-screen dial to match their representation of the target direction requiring them to concurrently maintain their representation of the target while planning and executing their response. Given that any target-aligned response in this task necessarily reflects the target direction, could the effect of priors be explained by response-related, rather than feature–specific, activity?

Participants were instructed to keep the mouse still until the response dial was presented. Therefore, decoding models trained during the observation period should be free from any influence of overt motor activity. We hypothesised, therefore, that if the effect of priors reflects modulation of motor activity, then it should be abolished when applying models trained only on stimulus-driven activity to unseen data recorded from the response period. To assess this possibility, we performed an exploratory temporal generalisation analysis using inverted encoding models trained between 250 and 750 ms after coherent motion onset, to recover motion information at every timepoint in held-out data. These time samples were selected for training since they comprised the approximate period over which robust decoding was observed at the group-level before motion offset (see **Figure 5B**). We then collapsed across the training timepoints to produce a single time-resolved measure of motion tuning for each level of prior precision (**Figure 7**). Consistent with the hypothesis outlined above, the effect of prior precision in the response period persisted, suggesting that enhanced neural tuning to the target direction under stronger priors reflects stimulus, rather than response-driven activity. Interestingly, feature-tuned information quickly decayed to chance levels under an uninformative prior, whereas similar activity under informative priors diminished more slowly.

**Figure 7.**
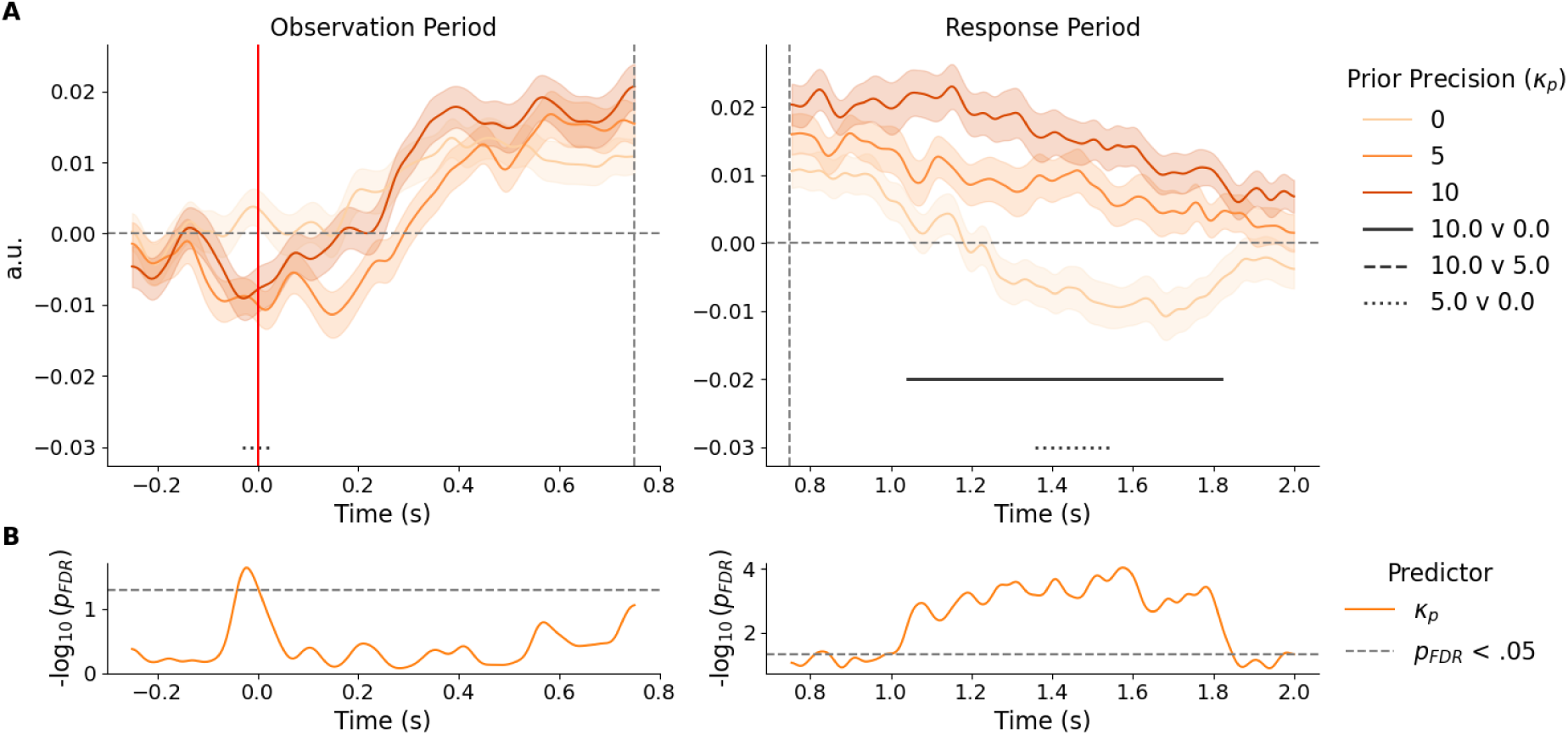
A) Decoding models trained during coherent motion observation reproduce the effect of prior precision when cross-generalised to the response period. Vertical red lines and grey dashed lines indicate coherent motion onset, and offset, respectively. Black solid, dashed and dotted lines indicate cluster significance p < .05 of pairwise condition comparisons. **B)** Log-transformed p values for fixed effect of precision of prior expectation on decoding accuracy. Horizontal dashes indicate the log-transformed significance threshold. For the purposes of presentation, each trace was smoothed using a temporal Gaussian filter (sigma = 4). All statistical comparisons were performed on the unsmoothed data.

## Discussion

Bayesian inference offers an elegant normative framework for describing perceptual decisions (Gold & Stocker, 2017; Ma et al., 2023). However, the degree to which neural representations of stimuli reflect these Bayesian computations remains unclear. Here, we focussed on the principle of precision-weighting – the notion that optimal decisions should combine priors and likelihoods, weighted by their respective precision – in a simple motion-estimation task. Using parametric manipulation of both likelihood and prior precision across two experiments, we found that behavioural reports demonstrate the hallmarks of precision-weighted inference: specifically, response accuracy reflected the combined precision of priors and sensory likelihoods, and the degree to which responses were biased toward the most likely stimulus values depended on the relative precisions of the likelihood and prior. These results are in line with previous empirical findings (Chalk et al., 2010; Körding & Wolpert, 2004; Rangelov et al., 2021) and suggest that the brain keeps track of task statistics and deploys this knowledge to optimise simple decisions in a statistically principled way (de Lange et al., 2018; Fiser et al., 2010; Summerfield & de Lange, 2014). But at what point/s are task priors integrated into the decision process?

Computational work using abstract accumulate-to-bound decision models has suggested that decision-biases most often result from changes to either the starting point or rate of evidence accumulation, each mechanism reducing the number of evidence samples necessary to trigger a decision in favour of the preferred outcome (Mulder et al., 2012; Ratcliff & McKoon, 2008; Summerfield & de Lange, 2014). A shift in the starting point for evidence accumulation may result from pre-existing evidence for stimulus values with a high prior probability relative to alternatives, whereas increasing the rate of accumulation implies an improvement in the quality of evidence being sampled, e.g., by sharpening perceptual representations when more probable stimuli are encountered. The continuous nature of our motion-estimation task does not lend itself to typical accumulate-to-bound sequential decision modelling, so instead we capitalised on the high temporal resolution of EEG to quantify the CPP, a well-established neural correlate of supramodal evidence-accumulation (O’Connell et al., 2012). Analysis of the CPP waveform as a function of stimulus precision revealed much faster ramping activity for high relative to low precision stimuli, replicating previous reports (Kelly & O’Connell, 2013; Rangelov et al., 2021; Rungratsameetaweemana et al., 2018; Stone et al., 2024). No clear gradation of ramping activity was observed when stratifying by the precision of prior expectations. Instead, stimuli drawn from an informative prior distribution elicited CPPs that peaked earlier, but with a lower overall amplitude relative to those drawn from a uniform distribution, suggesting less evidence was accumulated overall for stimuli with a higher prior probability. This result is consistent with recent work reported by Kelly et al. (Kelly et al., 2021), who found that validly cued motion directions elicited lower peak amplitude CPPs relative to neutral or invalid cues. It also supports the view that priors affect the starting point, but not the rate of evidence accumulation. (Interestingly, Kelly et al. (Kelly et al., 2021) also reported increased rates of evidence accumulation under valid probability cues, a result not borne out by our data.)

We next examined how neural representations of the target motion direction evolved over the course of a trial. Consistent with previous work from our group (McIntyre et al., 2022; Rangelov et al., 2021, 2024; Rangelov & Mattingley, 2020), feature-specific neural activity commenced shortly after target-onset and persisted well into the response-period after direct visual stimulation ceased. Interestingly, decoding accuracy improved in the response period compared with the observation period, suggesting that motion representations were strengthened as participants prepared their responses. Since aligning the response device with an internal representation of the target direction requires coordination between both sensory and motor processes, it is possible that feature-specific information is broadcast between systems leading to more robust overall representations during response production. This interpretation is broadly consistent with the widespread distribution of motion information at the beginning of the response period, though further investigation will be required to confirm this account.

Motion-decoding accuracy was sensitive to our manipulation of prior and likelihood precision; however, the time course of sensitivity was different for each factor. As expected, better decoding was achieved on high relative to low coherence trials, indicating that population-level stimulus representations encode the precision of the available sensory information (Rangelov et al., 2021). This difference persisted into the response period, indicating that information about stimulus reliability was readily available for response planning and execution. By contrast, decoding accuracy was only sensitive to the precision of the current prior during the response period, once motion information was no longer available to the participant. This finding runs counter to recent literature demonstrating early, and even pre-stimulus, effects of feature probability on representations decoded from M/EEG (Aitken et al., 2020; Kok et al., 2017; Tang et al., 2018). It is possible that this discrepancy can be traced to the comparatively weaker expectation for specific stimulus directions used in our study; participants here learned continuously-distributed priors over the space of possible motion directions, whereas previous work employed cues which probabilistically predicted one of only two (Kok et al., 2017) or five (Aitken et al., 2020) orientation/motion-directions with a high degree of certainty. Presenting fewer directions overall might have strengthened prior expectations for specific directions resulting in previously reported early effects of prior expectation. Nonetheless, our findings are in agreement with other studies that have challenged the extent to which recently learned prior probabilities can directly modulate early sensory processing (Bang & Rahnev, 2017; den Ouden et al., 2025; Feuerriegel, Vogels, et al., 2021; Hu et al., 2025; Rungratsameetaweemana et al., 2018; Rungratsameetaweemana & Serences, 2019).

Decoding accuracy was affected by the precision of prior expectations during the response period, with stronger priors driving more accurate decoding independent of the effect of stimulus precision (**Figure 6B**). This effect suggests that more probable motion directions were represented with greater fidelity, but only at later, response preparation stages. Since accurate estimation responses, by definition, will correlate strongly with the stimulus value, they necessarily elicit feature-selective patterns of neural activity which can be decoded the same way as activity patterns driven by the stimulus. One possibility is that more probable *actions* are represented with greater fidelity. On the other hand, it is possible that more probable *sensory features* drive heightened decoding accuracy in the response period. We tested this hypothesis with an exploratory temporal generalisation analysis in which decoding models trained to identify feature-selective activity patterns during observation of coherent motion were used to identify similar activity patterns during the response period. Because participants had to maintain fixation and not move the mouse until the response dial appeared, these decoders would not be sensitive to variance in activity patterns driven purely by response planning and execution. Instead, they should have been constrained by variance in neural signals associated only with accumulating evidence about the target motion direction. Intriguingly, the precision of prior expectations continued to affect neural representations in the response period: while decoding fell to chance levels in the uniform-prior condition, feature-specific information under informative priors was maintained, and numerically graded in its precision. This pattern of results suggests that the effect of priors in the main decoding analysis was, at least in part, driven by a pattern of activity that was already available during motion observation.

Under a Bayesian inference framework, sensory likelihoods and prior expectations are combined to produce a posterior belief whose mean reflects the precision-weighted average of each source of information. Given the behavioural results in the current study qualitatively followed this kind of inference scheme, with strong response biases toward the mean of the current task-prior, we originally expected neural representations of motion direction to be similarly biased. There was, however, no evidence of a systematic neural bias at any point during the trial, including during the response period, indicating that feature-specific activity does not track the Bayesian posterior or the eventual report, but instead faithfully reflects the veridical stimulus value. This result raises the important question of how behavioural biases arise if not from sensory representations. One possibility is that biases in continuous estimation tasks arise only during response preparation. Future investigations should consider using imaging methods with higher spatial resolution (e.g., MEG or fMRI) to specifically target response-selective brain areas.

Taken together, our results replicate a well-established finding in sensorimotor estimation that the brain learns new priors and uses them to guide decision-making in a precision-weighted manner consistent with Bayesian inference (Körding & Wolpert, 2004). However, in contrast to popular predictive processing accounts of perceptual decision-making (de Lange et al., 2018), we primarily observed effects of priors during response planning and execution rather than stimulus encoding.

## Acknowledgements

The authors wish to thank Douglas Cribb for assistance with data collection and Henry Beale for helpful advice on analysis. This work was supported by a grant from the Australian Research Council to JBM and DR (DP220104008). JBM was supported by an Australian National Health and Medical Research Council (NHMRC) Leadership (L3) Investigator Grant (GNT2010141).

