## Supplementary Figure 1 for "Late Integration of Prior Expectations During Precision-Weighted Perceptual Decisions"

### Supplementary Material

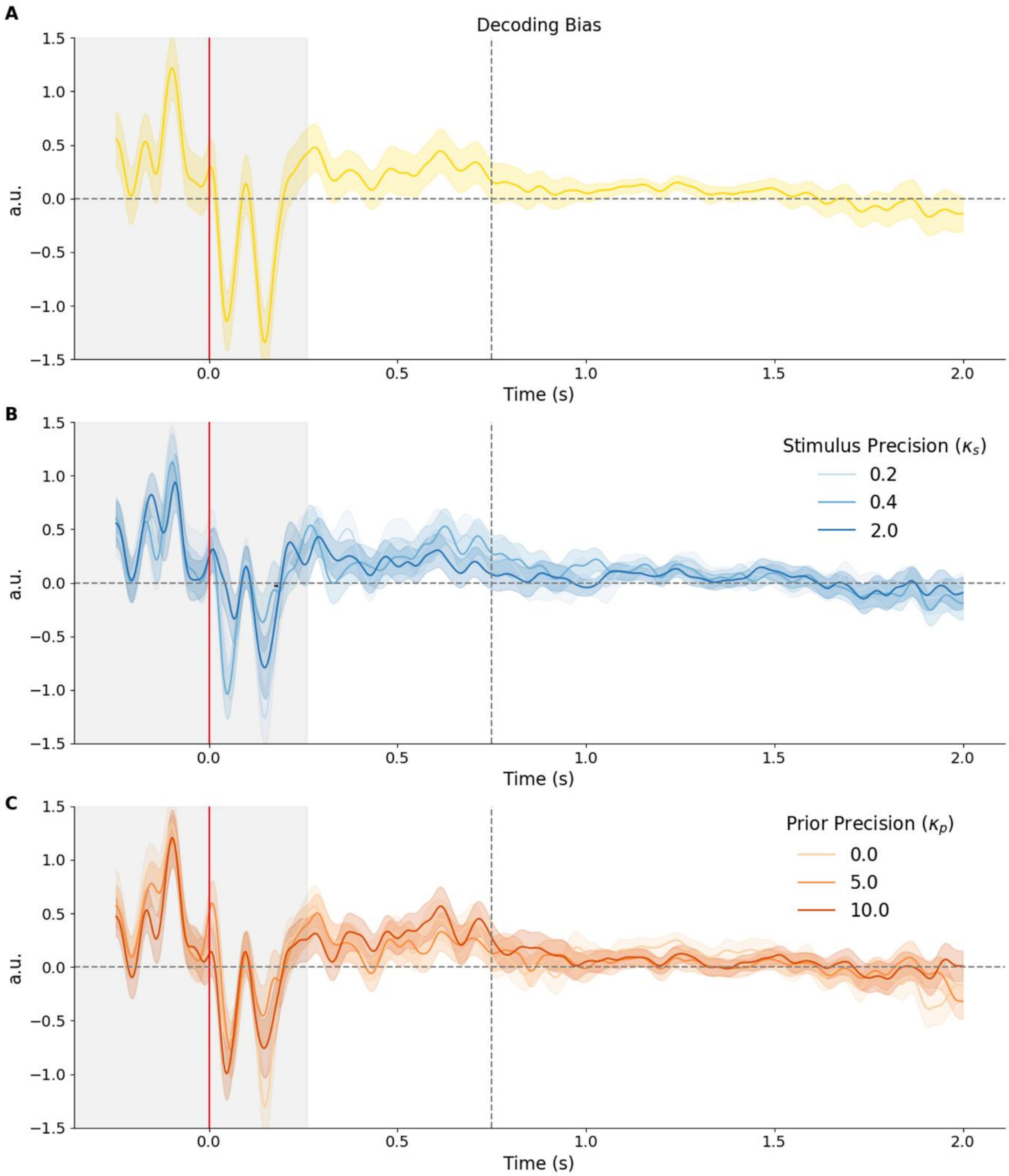


***Figure S1.*** *Time-resolved decoding bias expressed as the signed decoding error. Positive values indicate trial-wise decoded motion direction is biased towards the mean of the current prior.* ***A)*** *Grand decoding bias,* ***B)*** *decoding bias split by stimulus and* ***C)*** *prior precision. Colour-shaded regions indicate the between standard error in panel* ***A****, and the within-subjects standard error in panels* ***B*** *and* ***C****. The red and dashed vertical lines indicate motion signal onset and offset, respectively. Timepoints < 0.2s (shaded grey) are not interpretable since the calculation of bias uses a circular average of the decoding error that is undefined when decoding accuracy is close to zero.*
